# Microfluidic protein isolation and sample preparation for high-resolution cryo-EM

**DOI:** 10.1101/556068

**Authors:** Claudio Schmidli, Stefan Albiez, Luca Rima, Ricardo Righetto, Inayatulla Mohammed, Paolo Oliva, Lubomir Kovacik, Henning Stahlberg, Thomas Braun

## Abstract

High-resolution structural information is essential to understand protein function. Protein-structure determination needs a considerable amount of protein, which can be challenging to produce, often involving harsh and lengthy procedures. In contrast, the several thousands to a few million protein particles required for structure-determination by cryogenic electron microscopy (cryo-EM) can be provided by miniaturized systems. Here, we present a microfluidic method for the rapid isolation of a target protein and its direct preparation for cryo-EM. Less than 1 μL of cell lysate is required as starting material to solve the atomic structure of the untagged, endogenous *human* 20S proteasome. Our work paves the way for high-throughput structure determination of proteins from minimal amounts of cell lysate and opens new opportunities for the isolation of sensitive, endogenous protein complexes.

---

Knowledge of a protein’s architecture at high resolution is vital to understand its mechanics and chemistry. In recent years, cryogenic electron microscopy (cryo-EM) [1] has matured into a powerful method that can determine the architecture of biological macromolecules at the resolutions required to interpret the atomic fold of proteins [2, 3]. In the single-particle cryo-EM approach[4], an unsupported, thin layer of isolated protein complexes in amorphous (vitrified) ice is visualized at close to physiological conditions. Only several thousand to a few million imaged particles are needed to calculate a high-resolution, three-dimensional (3D) structure. Nevertheless, protein production, purification, and sample preparation for cryo-EM are nowadays considered the bottleneck for structure determination [5–7]. We have identified two dominating reasons for this: Firstly, significant amounts of protein must be produced. Conventional sample preparation for cryo-EM requires several microliters of a purified protein solution at a concentration of approx. 1 mg/mL per grid, from which extensive filter-paper blotting later removes the vast majority of protein particles [8, 9]. Secondly, both, protein purification and cryo-EM sample preparation are lengthy and harsh procedures. Mostly, high-yield expression systems are employed, and one or two chromatographic steps are needed to purify the protein particles. In addition, the classical cryo-EM sample preparation process that follows is a rough procedure [10], primarily because of the blotting step, and many proteins denature.

We recently developed a microfluidic cryo-EM grid preparation system termed *cryoWriter,* allowing the preparation of cryo-EM specimens from nanoliters of sample-solution [11–13]. Since the *cryoWriter* does not use paper blotting, it ensures that grid preparation is gentle and virtually lossless. Here, we report the combination of sample grid preparation using the *cryoWriter* with microfluidic protein purification [14], to determine the 3.5 Å cryo-EM structure of the untagged *human* 20S proteasome complex, which is the ‘catalytic core’ of the ubiquitin-proteasome system involved in 80 % of protein degradation [15] and an important drug target [16].

The microfluidic toolchain developed (Fig. 1A) consists of a module for affinity-isolation of the untagged protein from small quantities of cell lysate, followed by modules to write the purified protein onto a cryo-EM grid and vitrify the sample [12–14]. Briefly, antibody ‘fragment antigen binders’ (Fabs) are used to recognize and extract untagged target proteins from cell lysate. These Fabs are biotinylated with a photo-cleavable cross-linker, which binds with high affinity to the streptavidin functionalization of super-paramagnetic beads (Fig. 1B). First, the cell lysate is incubated with Fabs and super-paramagnetic beads. A <1 μL volume of this solution is then aspirated into the microcapillary of the *cryoWriter* system (Fig. 1C). The super-paramagnetic particles are immobilized in a ‘magnetic trap,’ isolating bound Fabs and their target proteins, while other cellular components are washed out. Ultra-violet (UV) light is used to break the photo-cleavable biotin cross-linker and the target proteins with the bound Fabs are eluted [14]. This procedure results in a 25 nL eluate, which is directly used to prepare one or more cryo-EM grids, while the magnetic particles are retained in the microcapillary.

**Figure 1:**
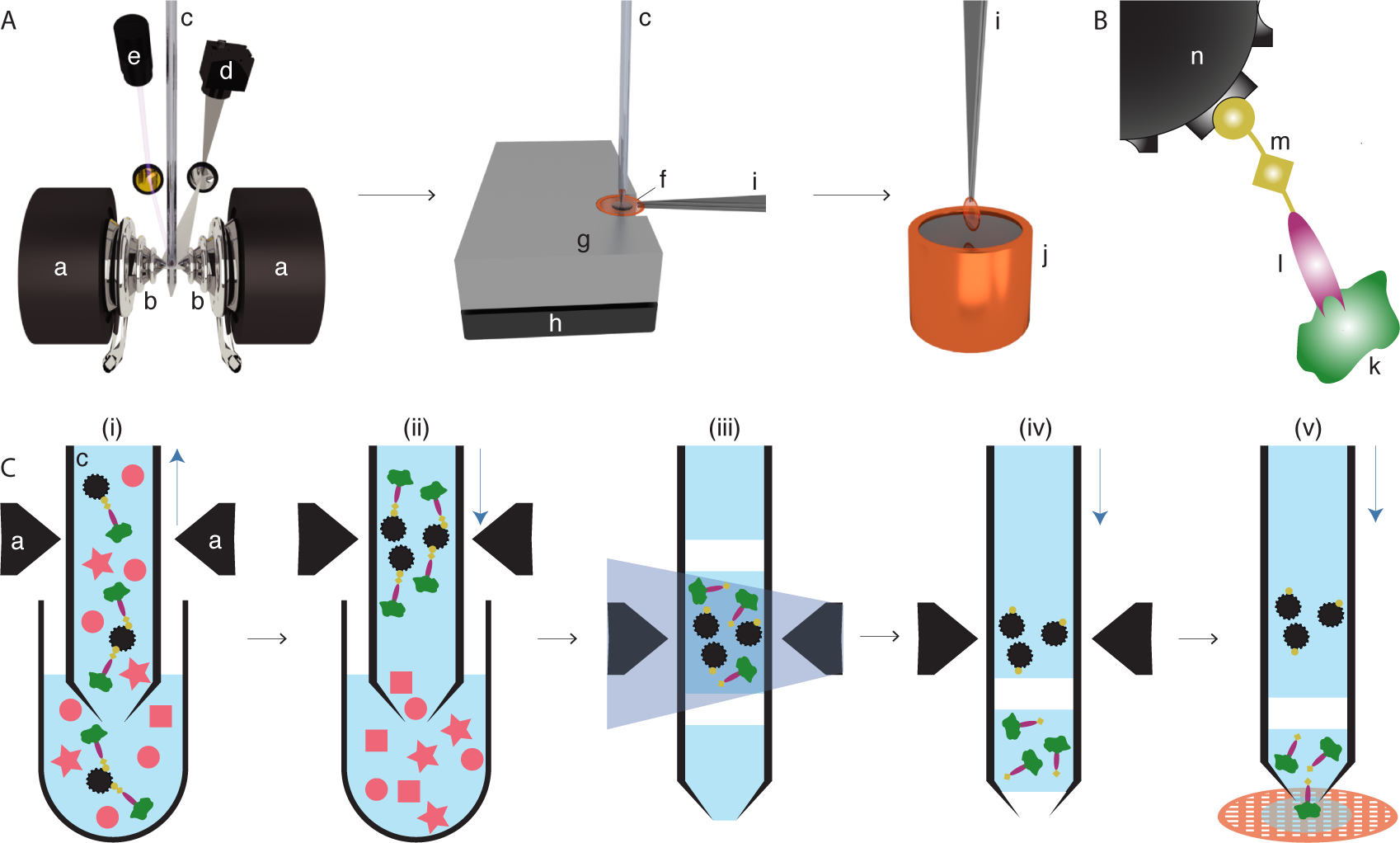
Schematic work-flow for microfluidic protein isolation and cryo-EM grid preparation. (**A**) Hardware for protein-isolation and cryo-EM grid preparation. The electromagnetic trap consists of two electromagnets (a) that produce a strong magnetic field gradient via their water-cooled iron tips (b). Sample processing in the capillary (c) is monitored by a camera (d), and a UV LED (e) allows photo-cleavage (see panel B) of the sample, both via mirrors. After protein isolation, the capillary nozzle is moved above a cryo-EM grid covered with a holey carbon film (f). The cryo-EM grid is positioned on a stage (g) that is temperature controlled by a Peltier element (h) and held with a Peltier-cooled tweezer (i). The isolated protein is directly written onto the grid and plunge-frozen in liquid ethane (j). (**B**) Composite material for ‘protein fishing.’ The target protein (k) is recognized by a Fab (l) that is covalently modified by a photo-cleavable cross-linker (m). The linker molecule ends with a biotin moiety, which strongly binds to the streptavidin coated bead (n). (**C**) Protein isolation work-flow. (i) Magnetic beads are incubated with biotinylated Fabs and cell lysate to capture the target structures (green). Less than 900 nL of sample is aspirated into the microcapillary for the protein isolation. (ii) The magnetic beads are immobilized in the magnetic trap (a). Non-bound lysate components (red) are flushed out. (iii) Illumination with UV light breaks the cross-linker. Before photoelution, two air bubbles are introduced and serve as boundaries to avoid dilution of the released proteins by diffusion (see Fig. SI 1 for details). (iv) Separation of the capturing magnetic beads and the eluted proteins. (v) The isolated target proteins are directly deposited on a cryo-EM grid for vitrification. The blue vertical arrows indicate the pump direction.

We used Fabs instead of full-length antibodies to avoid the cross-linking and aggregation that occurs when target proteins display more than one epitope. A linear cross-linker with three functional groups is used to connect the Fab to the super-paramagnetic beads. One end of the linker covalently attaches to the Fab while the biotin at the other end binds to the streptavidin-coated bead. A nitrobenzene moiety in between allows photo-cleavage of the linker upon illumination with UV light at 365 nm (Fig. 1B). Preparation of Fab fragments from antibodies and biotinylation with the cross-linker took ≈7 h; the biotintilated Fabs could be stored in the dark for several weeks at 4 °C.

The cell lysate was incubated with the Fabs and beads outside of the microcapillary in a 5 μL well. The cytosol/Fab mixture was incubated for 5 h, and then for an additional 1 h together with the super-paramagnetic beads. The use of miniaturized sample wells rather than the microcapillary, allowed several of these time consuming incubations to be carried out in parallel.

After incubation, a volume of 900 nL was aspirated from the well into the microcapillary, which was then precisely positioned between the iron tips of the particle trap system. In this setup, the trap is formed by electromagnets with water-cooled (4 °C) iron tips that concentrate the magnetic flux between the opposite poles and generate the field gradients required to trap the super-paramagnetic particles (see also supplementary information SI 1). The cooling system prevents heating and denaturation of the sample. A camera and magnifying lens system allow the procedure to be monitored. The 1 μm super-paramagnetic beads employed, can be easily immobilized in the magnetic trap and only marginally scatter photons with 365 nm wavelength, allowing efficient photoelution in 15 min [14]. Photoelution releases less non-specifically bound protein than the competitive elution typically used in other methods, because it does not change the buffer composition [17]. Just before elution, the beads were enclosed by two 6 nL air bubbles in a 25 nL buffer-plug, to prevent diffusion and Taylor dispersion [18]. After photoelution, the eluate was separated from the immobilized beads by the pump system and moved towards the apex of the microcapillary.

For cryo-EM specimen preparation, a sample carrier (grid) covered by a holey carbon film was placed on a temperature-controlled stage regulated to a temperature 7 °C above the environmental dew-point. This temperature offset builds a micro-environment on the grid surface, allowing controlled evaporation of the sample liquid. In a first step, 20 nL of the eluate containing the purified protein was written onto the grid, covering an area of ≈0.75 mm^2^. Excess sample was re-aspirated into the microcapillary, leaving a thin layer of the protein on the holey carbon film. This layer was left to settle for ≈50 ms with the grid still on the dew-point stage. The gentle evaporation that occurs during this time stabilizes [19] and thins the sample film. Finally, the written grid was rapidly removed from the stage and plunge-frozen, resulting in a vitrified ice layer [12,13]. In principle, several cryo-EM grids can be written with one 25 nL eluate. Interestingly, when this was done the first cryo-EM grid contained more protein particles than later grids. For the analysis presented here, we only used one cryo-EM grid, and this was the first grid prepared from the eluate. The whole *cryoWriter* process starting from protein isolation and ending with cryo-EM grid preparation could be performed in less than 1 h.

We employed the *cryoWriter* toolchain to isolate endogenous and untagged *human* 20S proteasome from commercially obtained HeLa cell lysate using Fabs generated from an antibody against the *α*4 subunit of the protein complex. As a positive control for the cryo-EM grid preparation, tobacco mosaic virus (TMV) particles were added to the elution buffer. The cryo-EM grid showed homogeneous ice layers (Fig. 2A&B) with the protein embedded in thin, vitreous ice. TMV particles and randomly oriented 20S proteasomes are visible, as well as smaller protein particles that are most probably unbound Fabs.

**Figure 2:**
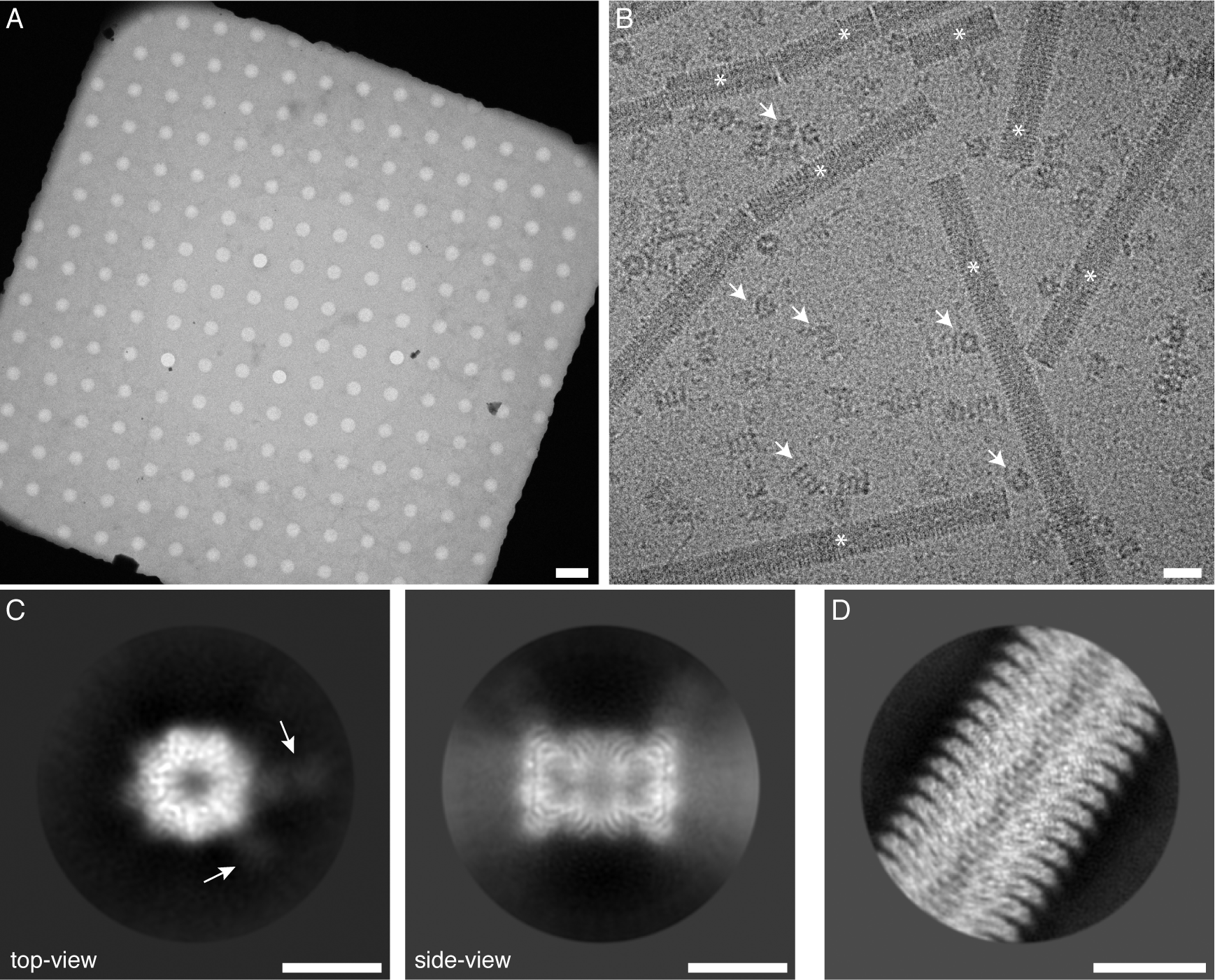
Sample quality and data-collection. (**A**) Overview image of a grid square and the holey carbon film, showing a thin film of vitreous ice. Scale bar: 4 μm. (**B**) High magnification image of the isolated 20S proteasome sample. Arrows indicate 20S proteasome as top and side views; asterisks denote TMV. Furthermore, small particles are visible in the background, most probably unbound Fabs. Scale bar: 20 nm. TMV was added to the elution buffer as positive control for the cryo-EM grid quality (see SI 2). The contrast of the image was increased using a Gaussian blur and subsequent histogram adjustment. (**C**) Selected projection averages of the 20S proteasome. Arrows indicate two bound Fabs recognizing the *α*4 subunit. Scale bar: 10 nm. (**D**) Typical projection average of TMV from the same cryo-EM grid as the 20S proteasome. Scale bar: 10 nm.

From only one grid, 523 dose-fractionated image stacks (movies) were recorded, yielding a total of 55 135 particles, which were processed with cryoSPARC v2 (Structura Biotechnology Inc.) [20] and RELION 3 [21] for structure determination. Fig. 2C shows typical projection class averages obtained from the 20S proteasome. A seven-fold pseudo symmetry is observed in the top view, with weakly visible Fab fragments attached to the two *α*4 subunits. The side-view exhibits the distinctive stack of *α-β-β-α* rings. Fig. 2D shows a projection class average of TMV. The secondary structure was visible in all projection-averages, demonstrating the excellent quality of the cryo-EM grid.

The 3D reconstruction of the human 20S proteasome shown in Fig. 3 has a resolution of 3.5 Å (for details see SI 2). The structure exhibits the typical dimeric, C2-symmetric arrangement of an *α*-*β* and a *β*-*α* ring-pair, each ring with a pseudo-7-fold arrangement of the respective subunits. We refined the X-ray structure [22] into our density map. The two Fabs against the *α*4-subunits aided the assignment of the individual components. Fig. 3B shows the *α* and *β* rings with all 14 models fitting the experimental densities in good agreement (see also Fig. 3 C & D, and SI 4). The densities of subunits *β*4, 5 and 6, are less well resolved (see also SI 3), which can be attributed to their catalytic activity [23]; we assume that the lower resolution reflects the greater flexibility they require to perform their biological function.

**Figure 3:**
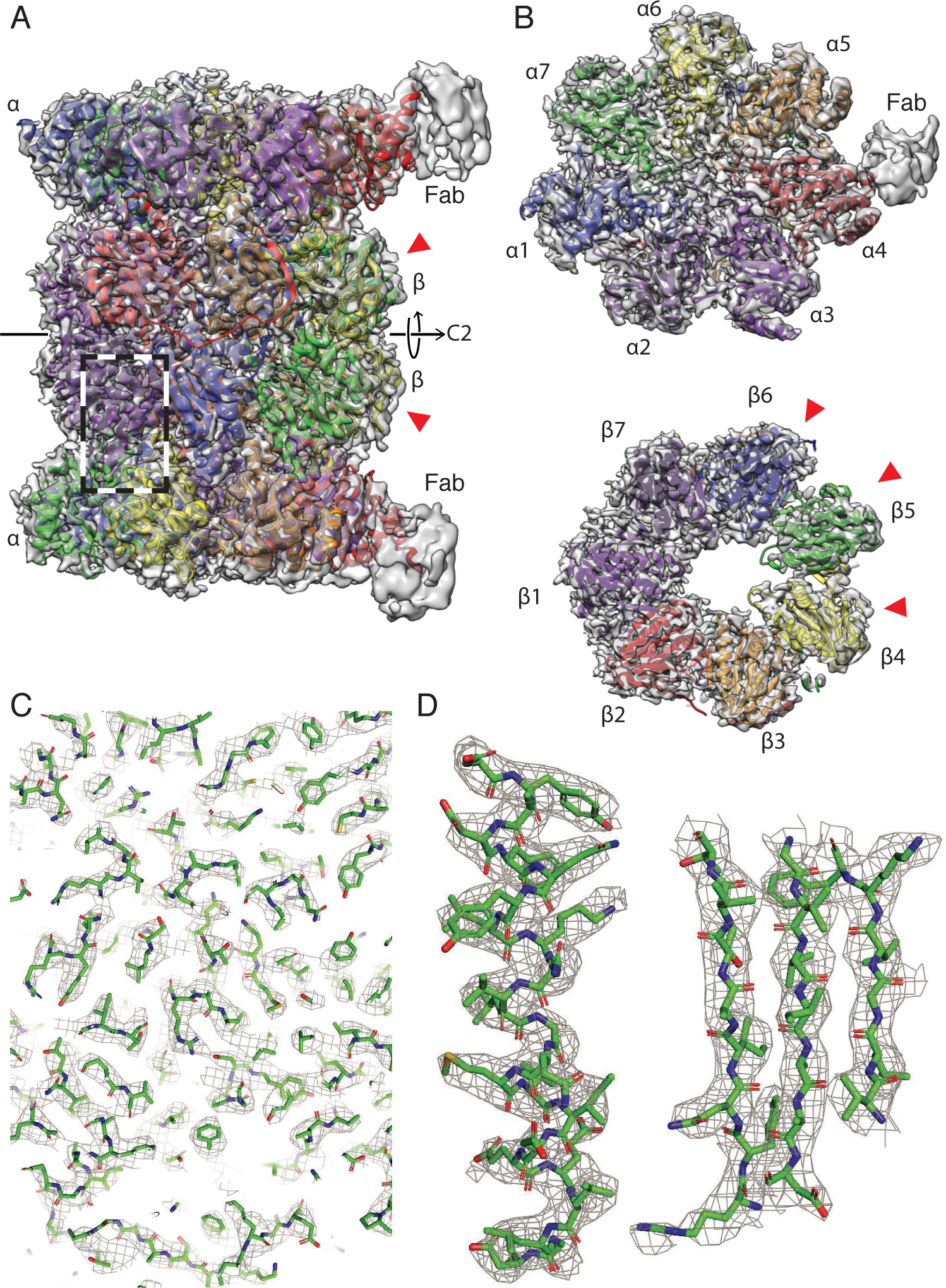
3D reconstruction of the human 20S proteasome. The red arrowheads indicate the catalytically active subunits. (**A**) Side view showing the two *α* and two *β* rings, all 14 subunits are fit into the mass-densities. Parts of the two bound Fabs are visible at lower resolution, due to the high flexibility of the attached Fabs. The C2 symmetry-axis is indicated. (**B**) Top view of an *α* and a *β* ring, both have a pseudo 7-fold symmetry. Different subunits are indicated by different colors. (**C**) A zoom into the side-view region indicated by the dashed box in panel A, documenting the quality of the data and model fitting. An enlarged view of one *a* helix and three strands of a *β*-sheet are shown in (**D**) as an example.

The reconstruction of the added TMV particles has a resolution of 1.9 Å (see SI 2 & 5), indicating that the microfluidic sample preparation did not limit the resolution of the *human* 20S proteasome. Unlike the archaeal T20S proteasome, which is often used as a cryo-EM test-sample thanks to its 14-fold D7 symmetry, the *human* 20S proteasome is only two-fold symmetric. The lower symmetry reduces the internal averaging by a factor of 7. Moreover, the similar subunits around the ‘pseudo-sevenfold’ axis can lead to misalignments of the particle projections. Together, these factors explain why existing cryo-EM maps are at the lower resolution of 3.5 Å [23, 24].

Several methods have been presented to improve sample preparation for cryo-EM, all having their own specific purpose. Examples include high-throughput grid preparation [25–27], time-resolved EM [28], and single-cell visual proteomics [11, 12]. Furthermore, antibody-functionalized EM-grids were proposed to ‘fish’ target proteins [29, 30], and affinity grids designed to capture proteins with engineered his-tags [31]. In both cases, the vitrified protein is supported by a continuous carbon film, which can limit the resolution obtained. Here, we combined microfluidics for protein isolation and purification with cryo-EM grid preparation, resulting in free-standing layers of vitrified protein samples that allow high resolution cryo-EM.

Microfluidic methods could potentially overcome the current bottleneck in cryo-EM, resulting from the large amount of protein used for sample preparation and the long and harsh conditions used for protein purification. We demonstrated that microfluidics (i) can deliver cryo-EM grids of high quality allowing <2 Å resolution from 20 nL of a sample and (ii) allows the isolation of endogenous proteins from less than 1 μL cell-lysate with subsequent structure determination at 3.5 Å. In addition, the method presented here significantly reduces both the amount of starting material and the time needed for the structural analysis of proteins, avoids harsh protein purification conditions, and eliminates the stringent and wasteful blotting steps otherwise employed on cryo-EM grid preparation.

The high efficiency of cryo-EM combined with the microfluidic approach will allow the structure of proteins that cannot be produced in large quantities to be studied. Up to now, structural studies were only possible if the protein could be over-expressed or produced in a large numbers of cells. The small sample volumes used during the microfluidic purification process allow a high protein concentration to be maintained, which is helpful if protein complexes fall apart upon dilution. Furthermore, the small sample volume also allows for ‘buffer conditioning’ [11], e.g., for the introduction of small ligands before sample vitrification. In addition, further optimization of the purification step and use of the *cryoWriter,* will almost certainly make it possible to perform the whole preparation in less than 1 h, facilitating the investigation of sensitive proteins targets. Finally, the toolchain presented here can also be combined with *in vitro* translation systems, which would enable high-throughput structure determination, or with a single cell lysis device [11,12, 32], which would bring the structural analysis of proteins originating from a single cell within reach.

## Acknowledgments

We thank the workshop of the Biozentrum of the University Basel for technical support, A. Fecteau-LeFebvre, D. Caujolle-Bert and K. Goldie for technical assistance, and A. Engel for discussions (all C-CINA, University of Basel). RELION 3 processing was performed at sciCORE (http://scicore.unibas.ch/) scientific computing center at University of Basel. TMV was kindly provided by R. Diaz-Avalos (now at Nanoimaging Services). We thank our former coworker Shirley Müller for critically reading the manuscript. **Funding:** We acknowledge support by the Swiss Nanoscience Institute (project P1401 & ARGOVIA MiPIS) and the Swiss National Science Foundation (projects 200021_162521 and 205320_166164), and the Swiss CTI (project 18272.1). **Author contributions:** C..S. performed microfluidic purification and grid preparation. S. A. and R. R. determined the cryo-EM structures. C. S. integrated the trap setup into the CryoWriter platform with the support of L. R. and P. O.. L. K. assisted with Titan Krios data collection. I. M. helped C. S. with model building. H. S. provided support and expertise in cryo-EM. T. B. conceived and coordinated the project. C. S., S. A., H. S., and T. B. wrote the manuscript, with backing from all authors. **Competing interests:** T. B. and H. S. declare the following competing financial interest: The cryoWriter concept is part of patent application PCT/EP2015/065398. **Data and material availability:** Image data are available at the EMPIAR database under EMPIAR-XXXX. The reconstructed volumes are available at the EMDB under EMDB-XXXX. The atomic coordinates are available at the PDB under PDB-XXXX.

## Material and Methods

### Generation of Fabs

Fabs were generated from an antibody against the *α*4-subunit of the human 20S proteasome (Enzo Life Sciences, BML-PW8120, Switzerland), using a commercial kit (Thermo Fisher Scientific Inc., # 44685, Switzerland) and following the provided protocol. After generation of Fabs, the buffer was changed to phosphate-buffered saline (PBS, pH 7.4, 150 mM NaCl, 1.5 mM KH_2_PO_4_, 8.1 mM Na_2_HPO_4_,2.6 mM KCl; Sigma, # D8537, Switzerland), and the Fabs were concentrated to a concentration of 0.28 mg/mL using a centrifugal filter unit (Sigma, # UFC501024, Switzerland).

### Biotinylation of Fabs

To biotinylatilate the Fabs, a six-fold molar excess of photo-cleavable NHS-biotin cross-linker (Fisher Scientific, # NC1042-383, USA) was added to the Fabs (0.28 mg/mL, 5.6 mM) and incubated at room temperature for 1 h. Excess, unbound cross-linker was removed using a spin desalting-column (Thermo Fisher Inc., # 87764, Switzerland) following the instructions from the kit-protocol.

### Binding of the 20S proteasome to magnetic particles

To ‘fish’ the human 20S proteasome, HeLa cell lysate (human origin, Enzo Life Sciences, BML-SW8750, Switzerland) was incubated in the presence of 0.7 mM of biotinylated Fabs at 4 °C for 5 h. Subsequently, super-paramagnetic Dynabeads™ (Thermo Fisher Scientific Inc., # 65602, Switzerland) were added to achieve a final concentration of 5 pM and incubated additionally for 1 h at 4 °C. Glycerol and ATP which maintain proteasome stability were intentionally not added to the buffers to ensure that the regulatory complexes dissociated from the 20S proteasome core complex [1].

### Extraction and vitrification of 20S proteasome

After the incubation step, a 0.3 μL volume of the cell lysate/Fabs/super-paramagnetic beads mixture was aspirated into the microcapillary of the *cryoWriter* setup at a flow rate of 0.4 μL/min, and passed through the electromagnetic trap (see Suppl. Info, Chapter SI 1). The beads were retained by the magnetic field gradient to form a bead plug. Subsequently, the flow direction was inverted and the particle plug was washed with 4 μL washing buffer (25 mM HEPES—KOH, pH 7.5, 5 mM MgCl_2_) at a flow-rate of 3 μL/min. The aspiration-wash cycle was repeated twice. In all, a total of 0.9 μL cell lysate was loaded into the microcapillary. At the end, the capillary was washed further with 45 μL of washing buffer at a flow-rate of 3 μL/min.

Before UV cleavage, the sample plug was enclosed between two 6 nL air bubbles that were introduced from the nozzle tip, and were separated by a volume of 25 nL washing buffer containing 1 mg/mL of tobacco mosaic virus (TMV) (Fig. 4). TMV was kindly supplied by Ruben Diaz-Avalos, Howard Hughes Medical Institute, USA. The hydrophilicity of the inner surface of the microfluidic capillary retains a thin hydration layer, which allows magnetic beads with loaded samples to remain within the aqueous phase while the air bubbles are introduced. The sample plug was then exposed for 15 min to UV light emitted at a wavelength of 365 nm by a 190 mW UV LED (Thorlabs, # M365L2, Switzerland).

**Figure 4:**
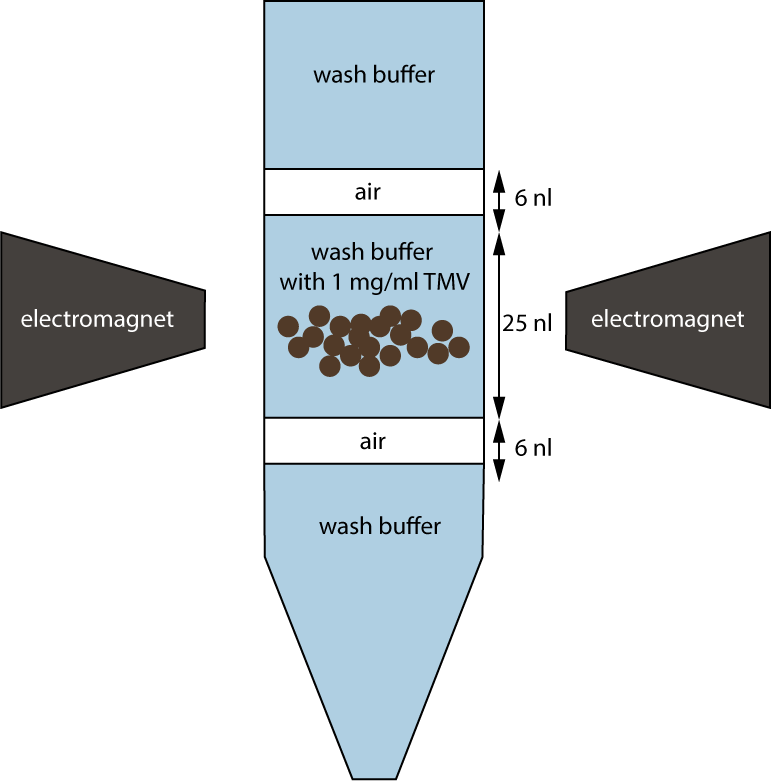
Illustration of the magnetic particle-plug enclosed between two 6 nL air bubbles for UV exposure and elution. The air bubbles act as barriers preventing diffusion and Taylor dispersion to keep the protein concentration as high as possible.

After photo-cleavage, 20 nL of the 25 nL eluate containing the purified protein was primed on a 400-mesh copper grid covered with a holey carbon film (R1.2/1.3, Quantifoil, Germany) that was glow discharged for 45 s in air plasma immediately before use. The cryo-EM grids were prepared as described [2], applying *protocol 1.* However, we adjusted the parameters for the sample: the stage temperature was 7 °C above the dew point temperature of the environment, *i.e.,* 10 °C on most days, and the settling time was 50 ms.

### Data Acquisition

Cryo-EM image data were collected on a FEI Titan Krios (ThermoFisher Scientific) transmission electron microscope, operated at 300 kV and equipped with a Gatan Quantum-LS imaging energy filter (GIF, 20 eV zero loss energy window; Gatan Inc.). Micrographs were acquired using a K2 Summit direct electron detector (Gatan Inc.) operated in dose fractionation mode (super-resolution 8k, 30 frames, 0.2 s/frames, 6 s total exposure) and controlled by the SerialEM [3] software. The physical pixel size was 0.812 Å and the total electron dose was 72 e^−^/Å^2^ per recorded image stack (movie). Micrographs were drift-corrected, dose-weighted, and Fourier-cropped to 4k, using MotionCor2 [4] via the FOCUS interface [5]. Additional data collection parameters are listed in Table 1.

**Table 1:** Data collection parameters.

| Data collection |  |
| --- | --- |
| Voltage | 300 kV |
| Physical pixel size | 0.812 Å |
| Super-resolution pixel size | 0.406 Å |
| GIF zero loss energy window | 20 eV |
| Defocus range | −1.0 μm to −2.2 μm |
| K2 operating mode | 8k super-resolution (with electron counting) |
| Number of frames | 30 |
| Frames per second | 0.2 |
| Total dose | $72 \text{ e}^-/\text{Å}^2$ |
| Number of micrographs | 523 |

### Image processing

An initial model for subsequent processing with RELION3 [6] was generated using cryoSPARC v2 (Structura Biotechnology Inc.) [7]. Particles were picked in cryoSPARC v2, and then 2D classified. An *ab-initio* model was generated using ≈19 000 particles selected from the best 2D classes, and subsequently refined to 4.1 Å resolution with C2 symmetry imposed.

For RELION3 processing, 55 135 particles were picked with Gautomatch (K. Zhang, www.mrc-lmb.cam.ac.uk/kzhang/) using templates projected from the map obtained using cryoSPARC v2, and low-pass filtered to 20 Å resolution. After 2D classification, 38 848 particles were 3D classified into 4 classes. The best class, containing 16015 particles was selected for 3D refinement. A generous mask derived from the refined map low-pass filtered to 15 Å and with a soft-edge of 6 voxels was used for postprocessing, yielding a resolution of 4.3 Å. Afterwards, rounds of CTF refinement including defocus refinement per-particle, astigmatism refinement per-micrograph, and beam-tilt refinement per-dataset were iterated with rounds of Bayesian polishing and 3D refinement. A final 3D refinement, using a softmask that excluded the flexible Fabs and employed solvent-flattened FSC curves to filter the 3D reference at every iteration, resulted in a map with a final global resolution of 3.5 Å after postprocessing (Fig. 7A). For more details on resolution estimation please see Suppl. Info., Chapter SI 2.

**Figure 7:**
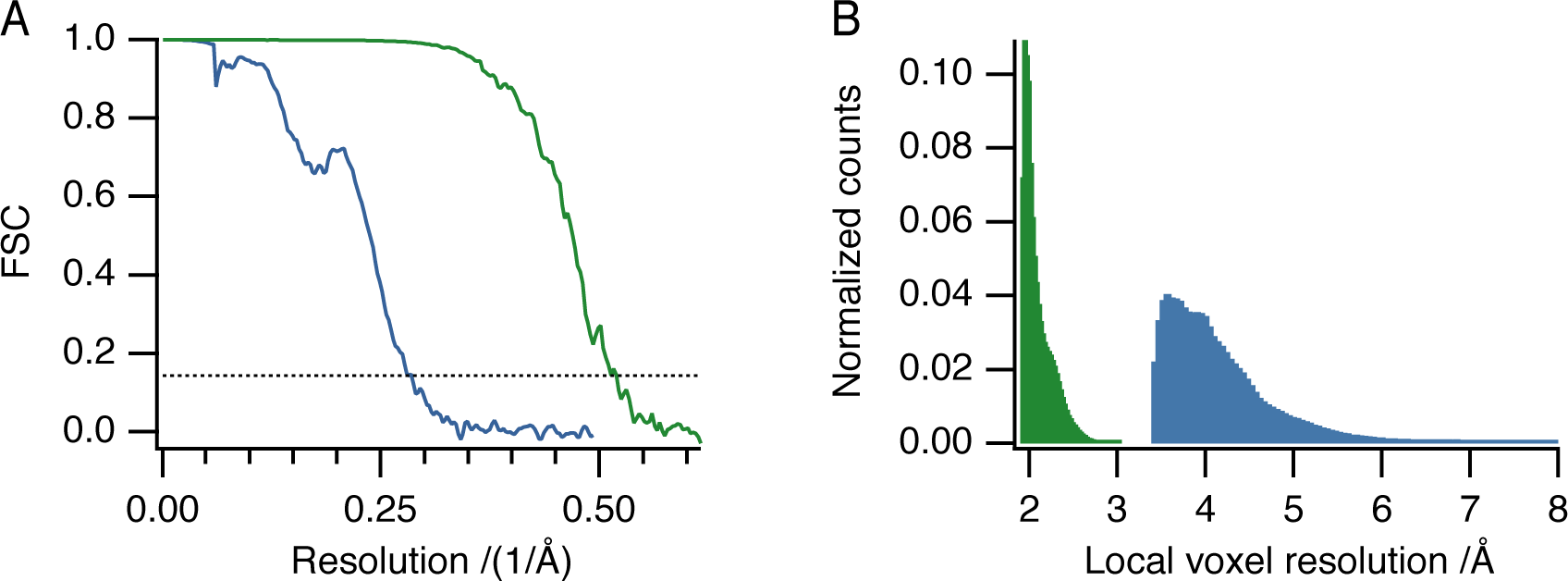
Resolution estimation for the 20S proteasome (blue) and TMV (green). (**A**) Fourier shell correlation from the RELION3 refinement. (**B**) Histograms depicting the normalized voxel counts per resolution range.

**Table 2:**
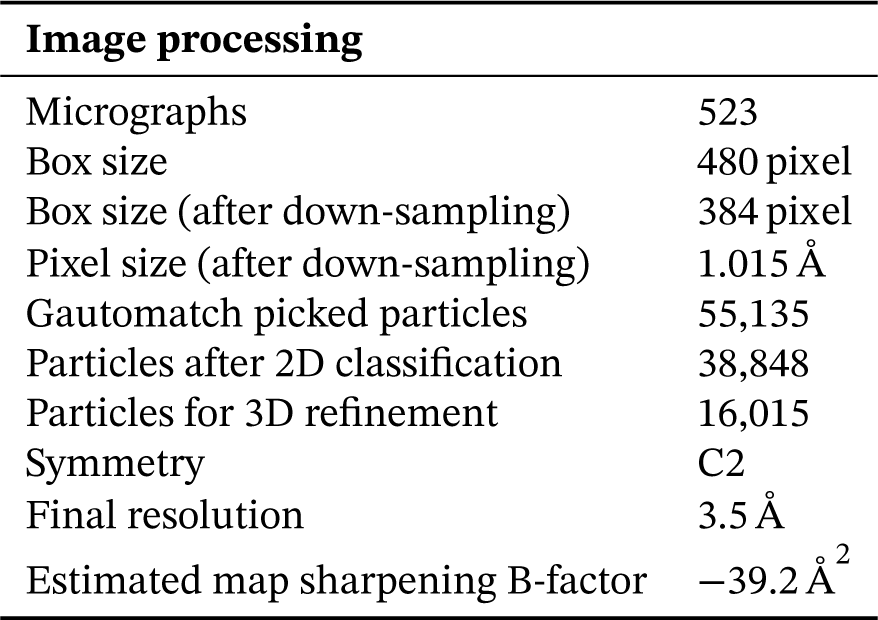
Image processing parameters used for structure determination with RELION3.

| Image processing |  |
| --- | --- |
| Micrographs | 523 |
| Box size | 480 pixel |
| Box size (after down-sampling) | 384 pixel |
| Pixel size (after down-sampling) | 1.015 Å |
| Gautomatch picked particles | 55,135 |
| Particles after 2D classification | 38,848 |
| Particles for 3D refinement | 16,015 |
| Symmetry | C2 |
| Final resolution | 3.5 Å |
| Estimated map sharpening B-factor | −39.2 Å<sup>2</sup> |

Helical processing of TMV was performed with RELION3 [6] from 481 micrographs of the same data set that contained 20S proteasome, following standard helical refinement and reconstruction procedures [8]. The final resolution of the TMV map was 1.9 Å (Fig. 7A).

### Model building

The model for the native human 20S proteasome was built based on the X-ray structure from Schrader et al. (PDB ID: 5LE5) [9]. Chimera [10] was used for the initial rigid-body fitting. The bound Fabs helped to identify the different subunits in the electron density map. For further processing, all heteroatoms were removed. Real space refinement with PHENIX [11] and manual adjustments in Coot [12] were done iteratively to obtain the final model, ending with a cycle of PHENIX real-space refinement. The graphics were generated using Chimera[10] and PyMOL[13].

## SI Supplementary information

### SI 1 Characteristics of the magnetic trap

The magnetic particle trap consists of two electromagnets arranged with opposite poles facing one another (Fig. 5). Attached iron tips extend the magnet cores. This concentrates the magnetic flux to a small area and forms a strong field gradient between the two tips (see Fig. 6). For magnetic particle trapping, the front part of the microcapillary segment just above the nozzle was placed in this region using a motorized XYZ linear translation stage, and the sample was aspirated. The magnetic force acting on the super-paramagnetic particles immobilized the beads. In experiments, super-paramagnetic particles with sizes from 15 nm to 1000 nm can be trapped and experience a magnetic force *F_m_* acting on a saturated particle according to:

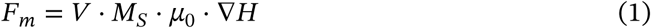

where V is the volume of the magnetic particles, *M*_s_ the saturation magnetization, *μ*_0_ the magnetic constant, and *H* the magnetic field strength. The current controls the field applied by the electromagnets. Due to the dependence on the volume (*V*), higher currents are needed in order to trap smaller particles. When the electromagnets are operated at maximum power, the core and the tips signihcantly heat up. For this reason, a cooling water system was implemented allowing the tips to be cooled to 4 °C by pumping water through a copper tubing twisted around them. Both, the copper tubing and the iron tips were coated with nickel to reduce corrosion. The magnetic trap is mounted on a motorized stage, allowing it to be moved in the vertical direction, independent of the microcapillary manipulator arm. For sample uptake, the magnetic trap and microcapillary were simultaneously moved down to insert the nozzle tip of the capillary into the sample well.

The parts used to build the magnetic trap are summarized in Table 3. Connecting parts were manufactured by the workshop of the Biozentrum, University of Basel, Switzerland. Plans are available upon request. Further cryoWriter parts used for grid preparation are described in [14].

**Figure 5:**
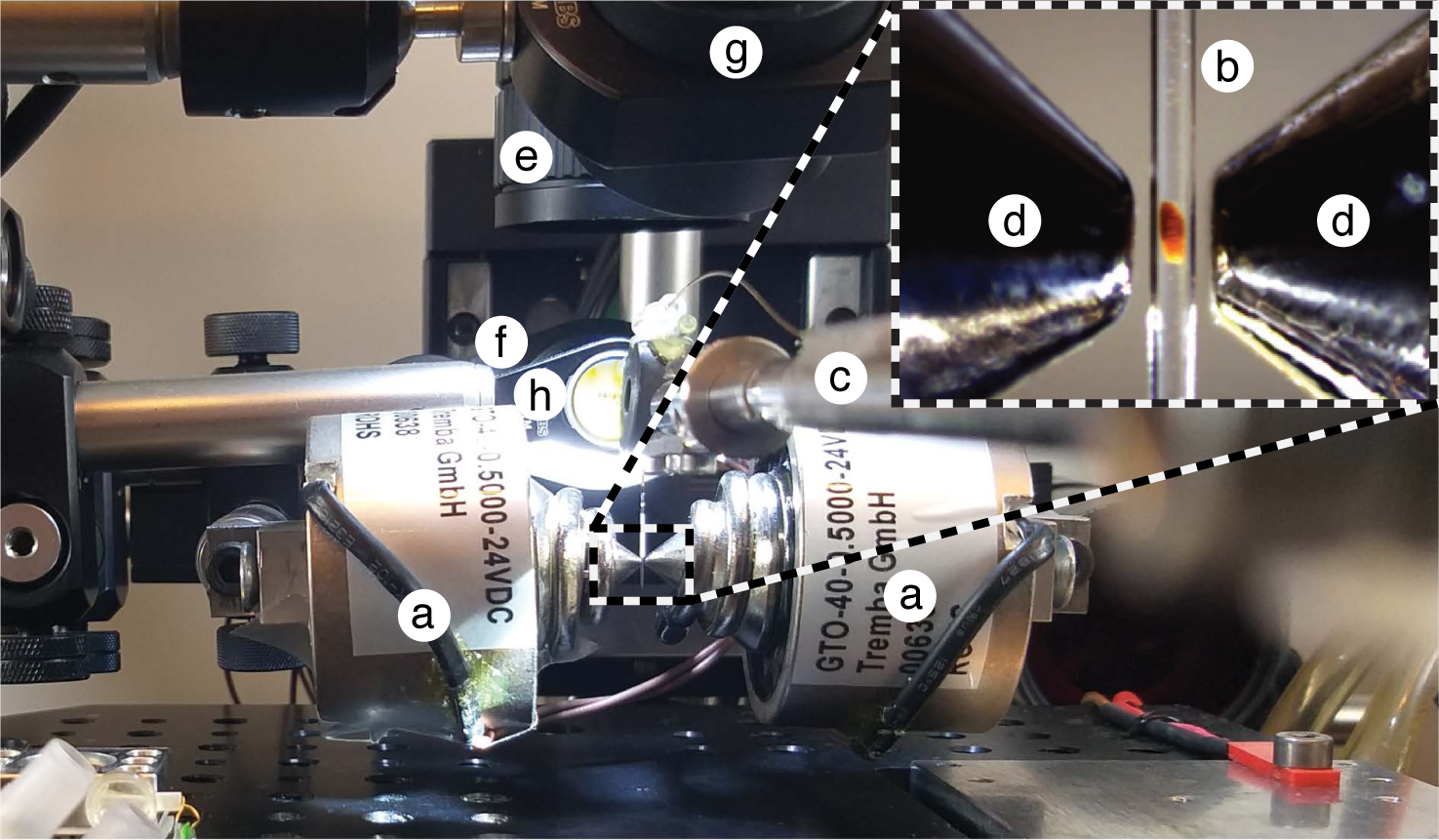
Image of magnetic the trap set-up. The opposite poles of two 4.8 W electromagnets. **(a)** are facing one another. A microcapillary **(b)**, held with the manipulator arm **(c)**, is placed between the two iron tips **(d)** of the electromagnets. The trapped magnetic plug can be observed with a camera **(e)** via a mirror **(f)**. The photo-cleavable cross-linker can be exposed to UV light using a LED **(g)** emitting light at 365 nm that is deflected by a mirror **(h)**.

**Figure 6:**
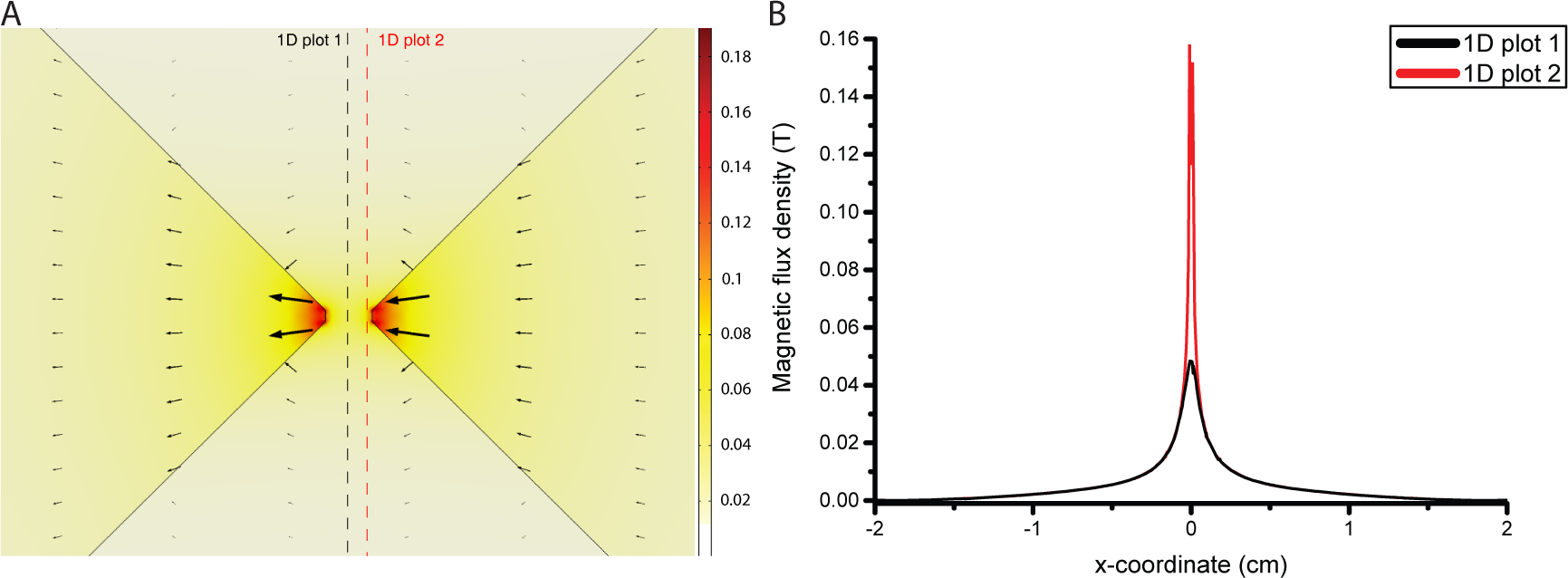
Finite element simulation of the magnetic field generated by the electromagnetic trap system. **(A)** 2D plot of the magnetic flux density around the iron tips (black solid lines). The direction of the magnetic fields is shown with black arrows. The size of the arrows is proportional to the field strength. The values in the color scale are given in Tesla (T). **(B)** Extracted 1D plots of the magnetic flux density from the 2D plot in (A) at the positions indicated with dashed lines. The figure shows that between the tips (x=0) the highest field gradients are observed. This leads to a magnetic force *F_m_* pushing magnetic particles towards this region.

**Table 3:** Parts list of the magnetic trap setup.

| Part Name | Part Number | Distributor |
| --- | --- | --- |
| Electromagnet | GTO-40-0.5000-24VDC | Conrad Electronic AG |
| Iron magnet tips with copper cooling tubes | - | In-house production |
| Cooling pump | 2449120 | Digitec Galaxus AG |
| Tubing for cooling system | 1025U04 | Parker Hannifin |
| Tubing for cooling system | 1025U06 | Parker Hannifin |
| Motorized stage for vertical trap movement | M-126.PD2 | Dyneos AG |
| Connector plate (trap-stage) | - | In-house production |
| Camera (Magnetic Trap) | NetFOculusFO124TC | NET GmbH |
| Optics (for magnetic trap camera) | MLM3X-MP | FRAMOS GmbH |
| Mirror (for magnetic trap camera) | PF07-03-P01 | Thorlabs |
| UV LED | M365LP1c | Thorlabs |
| Dielectric Mirror (to deflect UV beam) | BB0511-E01 | Thorlabs |
| Syringe pump controller | A3921000093 | Cetoni |
| Syringe pump dosing unit | A3921000095 | Cetoni |
| Liquid handling syringe | HA-80901 | BGB Analytik AG |
| Microcapillary tubing | TSP-150375-D-10 | BGB Analytik AG |
| Liquid handling nozzle | FS360-150-30-N-5-C12 | MS Wil GmbH |
| PressFit Connector (nozzle-microcapillary) | 2525LD | BGB Analytik AG |

Fig. 6 shows a finite element simulation of the magnetic flux density within the magnetic trap system, which was caluclated with COMSOL Multiphysics (COMSOL AB, Germany). Soft-iron material properties were applied to the magnet cores and tips using a non-linear B-H curve that includes saturation effects. The windings were modelled with coil features.

### SI 2 Fourier shell correlation and local resolution histograms

We used the Fourier shell correlation method [15] between two independently refined half-maps (‘gold standard’) [16] to estimate the resolution of our 3D reconstructions (Fig. 7A). A threshold of 0.143 on the FSC curve was adopted to define the resolution limit of the reconstruction [17]. The FSC curves were corrected for artificial correlations introduced by masking [18]. We found a global resolution of 3.5 Å for the 20S proteasome and of 1.9 Å for the TMV. These estimates correlate well with the features observed in the maps (see Fig. 3 of the main text for the 20S proteasome and SI 5 for the TMV). The estimated resolution is further corroborated by the local resolution histograms shown in Fig. 7B, which display the normalized counts of voxels at each resolution. The resolution for the proteasome varies significantly; for more details, see Chapter SI 3. The local resolution was assessed based on the half-maps using the method described in [19], as implemented in RELION3, with a map sampling of 25 Å (Fig. 7B).

### SI 3 Local resolution map of the 20S proteasome

Fig. 8 shows the local resolution calculated for the 20S proteasome using RELION3. Note that the active domains (indicated by red arrowheads in Fig. 8) are less well resolved than the other domains, and are associated with the catalytic subunits of the *β*-ring. Interestingly, the resolution of the *α*-ring around the pseudo-seven-fold axis is quite heterogeneous, whereas the *β*-ring exhibits lower resolution around subunits *β*4 and *β*5. Since subunits *β*4, 5 and 6 are the catalytically active elements [20], the lower resolution of subunits *β*4 and *β5* suggests that these are trapped in various conformations during cryo-EM grid preparation.

**Figure 8:**
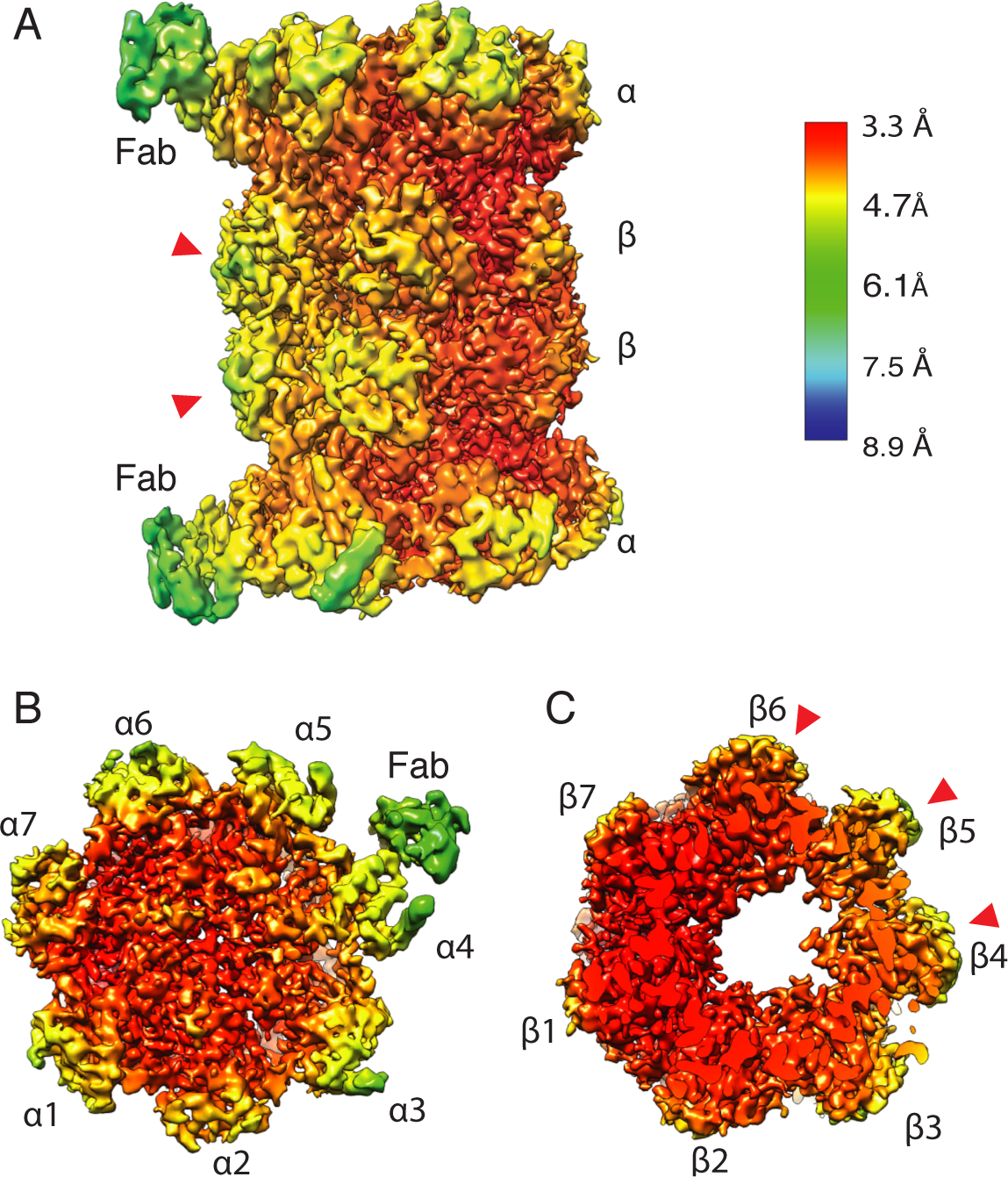
Local resolution map for the human 20S proteasome at a global resolution of 3.5 Å. (**A**) Side view with annotated Fabs. (**B**) Local resolution of the *α*-subunits. (**C**) Local resolution of the *β*-subunits. The red arrowheads indicate the catalytically active subunits.

### SI 4 Model building results

After initial rigid body fitting, the model was refined using PHENIX. The model was assessed by the ht to the experimental density and validated with clashscore, Ramachandran statistics, and good bond/angle lengths as shown in table 4.

**Table 4:** 20S proteasome model validation statistics.

| **Phenix Real-Space Refinement** |  |
| --- | --- |
| Map CC (around atoms) | 0.80 |
| **MolProbity[21]** |  |
| All-atom clashscore | 4.79 |
| Ramachandran favored | 96% |
| Ramachandran allowed | 2.43% |
| Ramachandran outliers | 0.26% |
| Rotamer outliers | 0.52% |
| Cb deviations | 0 |
| RMSD (bonds) | 0.008 |
| RMSD (angles) | 1.044 |

### SI 5 Three-dimensional map of TMV

Fig. 9 demonstrates the high quality of the microfluidic grid preparation. We report here the map of TMV at a global resolution of 1.9 Å. Compared to the 20S proteasome, this map has both higher global resolution (see FSC curve in SI 2, Fig. 7A) and less variable local resolution (Fig. 7B) due to the higher rigidity of the structure and the helical symmetry averaging imposed on the reconstruction.

**Figure 9:**
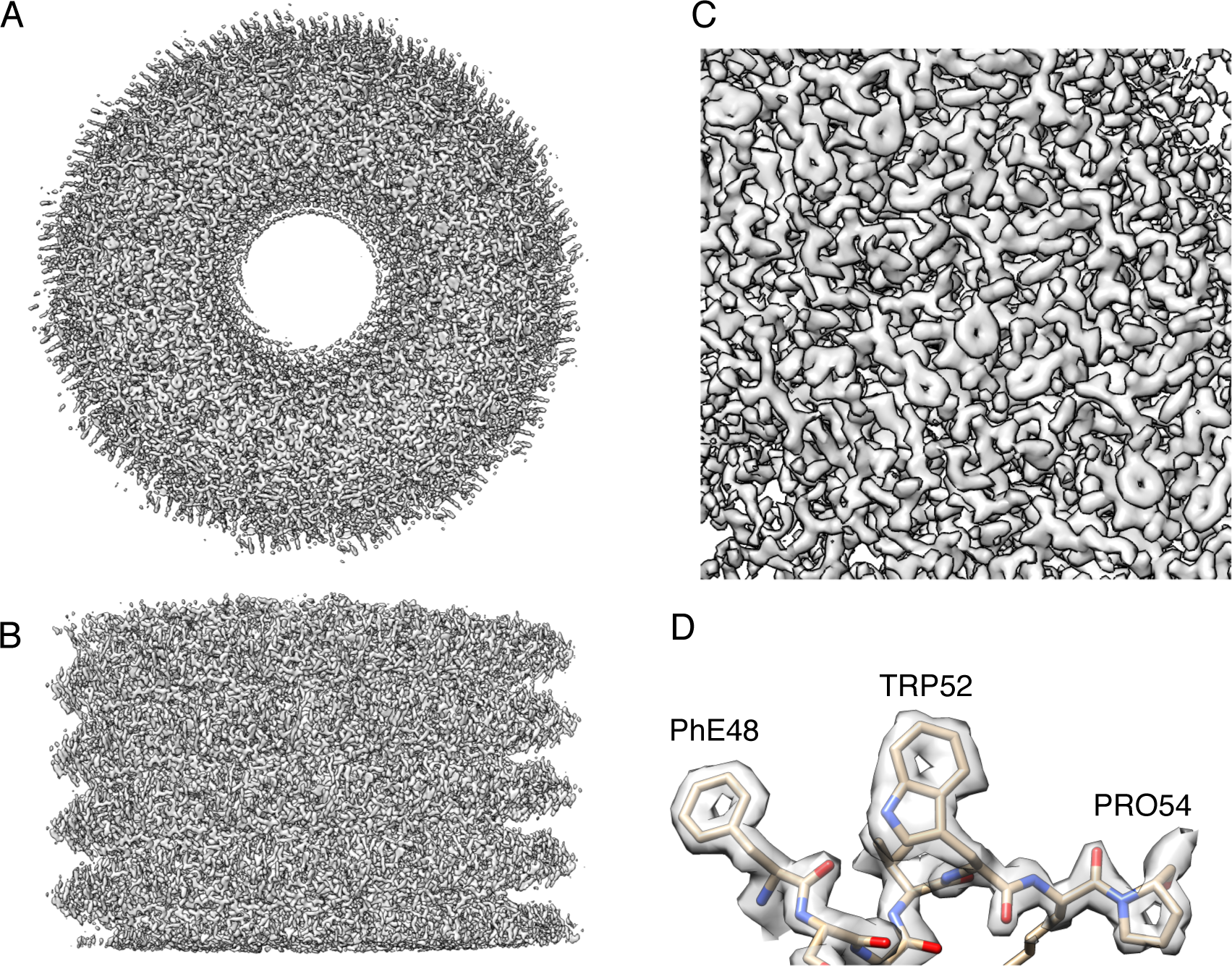
3D reconstruction of TMV at a global resolution of 1.9 Å showing the top view (**(A**) and the side view (**B**) of the electron density map. The zoom-in (**C**) demonstrates the high resolution features of the map. At this resolution it is possible to see the holes of the aromatic rings (**D**).

